# Hematology, biochemistry, and toxicology of wild hawksbill turtles (*Eretmochelys imbricata*) nesting in mangrove estuaries in the eastern Pacific Ocean

**DOI:** 10.1101/238956

**Authors:** Aubrey M. Tauer, Michael J. Liles, Sofía Chavarría, Melissa Valle, Sada Amaya, Gabriela Quijada, Oscar Meléndez, Stanley Rodríguez, Eric F. Lock, Ana V. Henríquez, Alexander R. Gaos, Jeffrey A. Seminoff

## Abstract

Sea turtles are a keystone species and are highly sensitive to changes in their environment, making them excellent environmental indicators. In light of environmental and climate changes, species are increasingly threatened by pollution, changes in ocean health, habitat alteration, and plastic ingestion. There may be additional health related threats and understanding these threats is key in directing future management and conservation efforts, particularly for severely reduced sea turtle populations. Hawksbill turtles (*Eretmochelys imbricata*) are critically endangered, with those in the eastern Pacific Ocean (Mexico–Peru) considered one of the most threatened sea turtle populations in the world. This study establishes baseline health parameters in hematology and blood biochemistry as well as tested for heavy metals and persitent organic pollutants in eastern Pacific hawksbills at a primary nesting colony located in a mangrove estuary. Whereas hematology and biochemistry results are consistent with healthy populations of other species of sea turtles, we identified differences in packed cell volume, heterophils and lympohcyte counts, and glucose when comparing our data to other adult hawksbill analysis (1), (2), (3). Our analysis of heavy metal contamination revealed a mean blood level of 0.245 ppm of arsenic, 0.045 ppm of lead, and 0.008 ppm of mercury. Blood levels of persistent organic pollutants were below the laboratory detection limit for all turtles. Our results suggest that differences in the feeding ecology of eastern Pacific hawksbills in mangrove estuaries may make them less likely to accumulate persistent organic pollutants and heavy metals in their blood. These baseline data on blood values in hawksbills nesting within a mangrove estuary in the eastern Pacific offer important guidance for health assessments of the species in the wild and in clinical rehabilitation facilities, and underscore the importance of preventing contamination from point and non-point sources in mangrove estuaries, which represent primary habitat to hawksbills and myriad other marine species in the eastern Pacific Ocean.

## Introduction

Disease can cause declines in wildlife populations, especially those that are already threatened or vulnerable (4) (5) (6) (7). Baseline hematology and biochemistry blood parameters are useful indicators for the assessment of the health status of wild nesting sea turtle populations (8) and are especially helpful in clinical rehabilitation facilities (9). However, reference ranges for hematology and blood biochemistry are not widely available, with many reported values derived from captive animals that may not be representative of wild individuals. Additionally, data from one population of a species often are used as references for other populations, despite potential within-species variation (10) (11) (12) (13).

Hawksbill turtles (*Eretmochelys imbricata*) exemplify a species whose life history may vary widely among populations in distinct ocean basins (14) (15). In the Atlantic and Indo-Pacific, adult hawksbills primarily inhabit coral reef ecosystems (16) (17) (18) and can embark on long-distance (>2,000 km), offshore migrations between nesting and foraging areas (e.g., (19) (20). Hawksbills in the eastern Pacific, however, often associate with mangrove ecosystems (21) (14) (22) (15) and undertake particularly short (<300 km) and neritic (<5 km) post-nesting migrations (23) (24). The marked difference in life history among hawksbills in these ocean basins could greatly influence general health parameters, which are largely unknown for adult hawksbills (3) and which have never been analyzed for individuals inhabititing mangrove estuaries. The availability of reference ranges is paramount for different populations of the same species and even subspecies, as values may even vary amongst a small population depending on diet and ecological variables (25).

Hawksbills are critically endangered globally according to the International Union for the Conservation of Nature’s (IUCN) Red List (26) and the population in the eastern Pacific is among the most endangered Regional Management Units (27) for sea turtles worldwide (28). Fewer than 700 adult female hawksbills are estimated to remain in the entire eastern Pacific Ocean (29) (15), where >80% of these individuals nest on beaches in mangrove estuaries of El Salvador and Nicaragua (30) (31) (15). These same mangrove ecosystems also provide important developmental habitat for juvenile and sub-adult hawksbills (14) (32). Known threats to this species in the region include incidental capture in coastal fisheries, human consumption of eggs, and alteration of nesting habitat (29) (31). An additional, albeit understudied potential threat to hawksbills inhabiting mangrove estuaries, is contamination by chemicals used in aquacultural and agricultural operations, including persistent pesticide residues from shrimp ponds (33) and toxic compounds dicharged by surrounding rivers (34). These contaminants have been documented as negatively influencing myriad species, including estuarine fish species (35) mollusks (36) and marine turtles (37). If these contaminants are present in mangrove estuaries, reliance on such habitats could have direct impacts on health of hawksbills.

In this study, we measured blood biochemistry, hematology, and toxicological parameters in wild adult female hawksbills nesting in the Bahía de Jiquilisco mangrove estuary complex in El Salvador to establish baseline health data for one of the most important hawksbill nesting areas in the eastern Pacific. This information will establish a baseline for these parameters and aid in long-term evaluation of the health status of this severely depleted population and serve to guide future management and conservation efforts, as well as to facilitate comparisons among hawksbill populations in other oceanic basins.

## Materials and Methods

### Study site

Bahía de Jiquilisco (13°13’N, 88°32’W) is located in the Department of Usulután on the south-central coast of El Salvador (Fig. 1), and is a National Conservation Area, RAMSAR wetland, and UNESCO Biosphere Reserve. It contains the largest mangrove forest in El Salvador (635 km^2^), and includes numerous islands, channels, and estuaries, with moderate development at some nesting beaches (31). Bahía de Jiquilisco has 42.1 km of hawksbill nesting habitat that includes eight discernable fine grained sand beaches with fragmented second growth coastal forest and fruit tree plantations adjacent to the high water line (15) which host ~40% of hawksbill nesting activity in the eastern Pacific (29) (31) (38).

Fig. 1. Locations of hawksbill nesting beaches with patrolled shoreline (black lines) at Bahía de Jiquilisco, El Salvador, 2013–2014.

### Beach Monitoring and Turtle Measurements

Hawksbill nesting occurs primarily between April and October, with a peak in June–July. We conducted beach patrols from 1 April to 15 October 2013–2014 at Bahía de Jiquilisco, where project personnel and an extensive network of >100 trained local egg collectors monitored nesting habitat from 18:00 to 06:00 daily by foot and boat in search of female hawksbills. We identified turtles by Inconel tags (Style 681, National Brand & Tag, Newport, KY, USA) located on the second proximal scale of both front flippers and internal passive integrated transponders (PIT tags; Biomark, Boise, ID, USA) in the right front flipper; Inconel and PIT tags were either present from application during previous tagging seasons or were applied after egg laying was completed (15). For each female hawksbill encountered, we measured curved carapace length (nuchal notch to posterior-most tip of marginal scutes; CCL) and in 2013 we performed a complete visual and physical examination, noting all epi-biota on the turtle and body condition.

### Sample Collection and Analyses

We collected up to 12 ml of blood from the dorsal cervical sinus using a 10 ml syringe and 18 gauge 1.5 inch needle and immediately transferred the sample into a red-top glass serum separator tube and sodium heparin vacutainer tubes. They were not refrigerated prior to processing. Blood smears were made in our field base camp from sodium heparin-treated blood and were fixed with 99% methanol on glass slides and air dried.

We initially processed the blood in the field within 6–8 hours of blood collection. Packed cell volumes were performed using a tabletop centrifuge and whole blood in sodium heparin tubes was transferred to 1 ml cryotubes and frozen in liquid nitrogen for heavy metal analysis. The remaining blood was spun for 10 minutes at 2000 RPMs and the serum separated and frozen in cryotubes in liquid nitrogen in the field, which were subsequently stored in −20° C freezers at the University of El Salvador. Samples collected in 2013 were shipped in dry ice to the United States for hematology, serum biochemistry, heavy metal, and toxicology analyses, whereas in 2014, plasma biochemistry analyses were conducted at Centro Scan (San Salvador, El Salvador). The results were pooled for determining biochemistry reference ranges after determining that there was no statistical difference between the two sample sets.

For hematology, blood films were stained at the Minnesota Zoo with DipQuick stain (Jorgenson Laboratories, Loveland, CO, USA) for manual differential accounts of circulating white blood cells and for hemo-parasite identification. Total white blood cell counts were estimated. Samples for serum biochemistry were shipped on dry ice for processeing at Marshfield Laboratories (Marshfield, WI, USA). The biochemical panel included alanine aminotransferase (ALT), aspartate aminotransferase (AST), alkaline phosphatase, cholesterol, CO2, creatine kinase (CK), glucose, lactate dehydrogenase (LDH), calcium, phosphorous, potassium, sodium, chloride, bicarbonate, total protein, anion gap and uric acid (UA). Reference intervals for biochemistry and hematology variables were computed using the package referenceIntervals for R (39) Parametric 95% reference intervals were computed, and the one-sample Kolmogorov-Smirnoff test (40) was used to assess the distributional assumption. For variables with a Kolmogorov-Smirnoff p-value less than 0.01, a non-parametric 95% reference interval was determined instead, with endpoints given by the 0.025 and 0.975 sample quantiles of the observed data.

Blood samples were screened at the California Animal Health and Food Safety Laboratory (San Bernadino, CA, USA) for heavy metals (arsenic [detection limit = 0.010 ppm], lead [0.050], and mercury [0.010]) and persistent organic pollutants (POP), including organochlorine insecticides (aldrin [0.010], alpha-BHC [0.010], gamma-chlordane [0.010], technical chlordane [0.050], pp-DDE [0.020], pp-DDD [0.020], pp-DDT [0.020], dicofol [0.020], op-DDE [0.020], op-DDD [0.020], op-DDT [0.020], dieldrin [0.010], endosulfan I [0.010], endosulfan II [0.010], endrin [0.010], HCB [0.010], heptachlor [0.010], heptachlor epoxide [0.010], lindane [0.010], methoxychlor [0.010]), mirex [0.010], toxaphene [0.400] and polychlorinated biphenyl (Arochlor 1221, 1232, 1242, 1248, 1254, 1260, 1262 [0.200 and 0.400]). Mean toxicity levels were determined, with 95% confidence intervals, for arsenic, lead, and mercury. If observations were missing below the limit of detection, the mean and standard deviation were inferred via maximum likelihood under the assumption that the data have a log-normal distribution that is left-censored below the limit of detection using the package censeReg for R (41). All data and analysis is publicly available as an annotated reproducible R code file at and https://doi.org/10.6084/m9.figshare.5702818 https://doi.org/10.6084/m9.figshare.5702779.

## Results

We encountered and examined 66 nesting hawksbills at Bahía de Jiquilisco in 2013–2014, which had a mean carapace length of 84.9 cm (SD 5.8, range = 71.0–96.6) and appeared in good general health. Physical exam findings in 2013 included two turtles that were covered in approximately 5% of epi-biotic growth; nearly all other individuals were less than 1%. Six individuals exhibited carapace damage, including missing scutes, although all appeared to have healed from the injuries. One individual had a fairly large deformity of her distal carapace, but was mobile, in good body condition, and did not have difficulty depositing eggs. Additionally, one individual had a small tumor on the right rear flipper, but logistical limitations prevented biopsy collection. Hematologic values are presented in Table 1. No hemo-parasites were observed for the 28 hawksbills evaluated in 2013. Table 2 provides the serum biochemistry reference ranges for blood collected in plain serum separator tubes in 2013 and blood plasma from sodium heparin tubes in 2014, including liver enzymes, total protein, electroyltes, and uric acid.

**Table 1.** Hematology reference intervals for wild hawksbill turtles (*Eretmochelys imbricata*) nesting at Bahía de Jiquilisco, El Salvador, 2013 (n = 28).

| Parameter<sup>a</sup> | 95% reference interval |  | KS p-value<sup>b</sup> |
| --- | --- | --- | --- |
|  | Low | High |  |
| PCV (%) | 23.0 | 33.6 | 0.7130 |
| WBC ($\times 10^3/\mu\text{L}$) | 1228.2 | 10,871.8 | 0.0528 |
| Heterophils (%) | 43.4 | 93.6 | 0.3306 |
| Heterophils ($\times 10^3/\mu\text{L}$) | 0 | 8549.7 | 0.2120 |
| Lymphocytes (%) | 41.8 | 42.6 | 0.8327 |
| Lymphocytes ($\times 10^3/\mu\text{L}$) | 87.5 | 2517.7 | 0.8282 |
| Monocytes (%) | 0.3 | 11.8 | 0.1358 |
| Monocytes ($\times 10^3/\mu\text{L}$) | 0 | 748.3 | 0.4461 |
| Basophils (%) | 0 | 7.1 | 0.5905 |
| Basophils ($\times 10^3/\mu\text{L}$) | 0 | 398.4 | 0.8808 |
| Eosinophils (%)<sup>c</sup> | 0 | 5 | <0.0001 |
| Eosinophils ($\times 10^3/\mu\text{L}$)<sup>c</sup> | 0 | 290 | <0.0001 |

<sup>a</sup>PCV, packed cell volume; WBC, white blood cells.
<sup>b</sup>KS p-value, Kolmogorov-Smirnoff p-value.
<sup>c</sup>Indicates non-parametric 95% reference interval.

**Table 2.** Serum chemistry reference intervals for wild hawksbill turtles (*Eretmochelys imbricata*) nesting at Bahía de Jiquilisco, El Salvador, 2013–2014 (n = 66).

| Parameter<sup>a</sup> | n | 95% reference interval |  | KS p-value<sup>b</sup> |
| --- | --- | --- | --- | --- |
|  |  | Low | High |  |
| Glucose (mg/dL) | 51 | 57.5 | 142.0 | 0.1771 |
| AST (U/L) | 51 | 18.3 | 74.3 | 0.8810 |
| ALT (U/L) | 33 | 16.3 | 78.1 | 0.8429 |
| ALP (U/L) | 51 | 25.7 | 93.6 | 0.4668 |
| CK (U/L)<sup>c</sup> | 51 | 121 | 1296.2 | <0.0001 |
| LDH (U/L)<sup>c</sup> | 51 | 135.6 | 1645.7 | 0.0009 |
| Cholesterol (mg/dL) | 51 | 107.1 | 366.8 | 0.8260 |
| TP (g/dL) | 51 | 2.6 | 5.0 | 0.6850 |
| Phosphorus (mg/dL) | 18 | 3.4 | 16.1 | 0.6876 |
| Calcium (mg/dL) | 36 | 1.0 | 21.2 | 0.0379 |
| Sodium (mmol/L) | 51 | 140.7 | 169.4 | 0.8217 |
| Potassium (mmol/L) | 18 | 3.7 | 5.7 | 0.9910 |
| Chloride (mmol/L) | 51 | 96.9 | 148.5 | 0.1106 |
| Bicarbonate (mmol/L) | 18 | 9.8 | 33.5 | 0.9538 |
| Uric Acid (mg/dL) | 33 | 1.0 | 1.8 | 0.1846 |
| Anion Gap (mmol/L) | 18 | 7.6 | 36.8 | 0.2089 |

<sup>a</sup>AST, aspartate aminotransferase; ALT, alanine aminotransferase; ALP, alkaline phosphatase; CK, creatine kinase; LDH, lactate dehydrogenase; TP, total protein.
<sup>b</sup>KS p-value, Kolmogorov-Smirnoff p-value.
<sup>c</sup>Indicates non-parametric 95% reference interval.

Levels of arsenic, lead, and mercury are presented in Table 3. Arsenic had the highest level, with a mean of 0.245 ppm (95% confidence interval = (0.10, 0.39)). Arsenic was detectable in all of the samples collected (n = 28). Lead and mercury had lower mean levels of 0.045 (95% confidence interval = (0.038, 0.056)) and 0.008 (95% confidence interval = (0.004, 0.017)) respectively. Samples from all turtles tested for POPs (n = 28) were below the detectable limits.

**Table 3.** Heavy metal blood values for wild hawksbill turtles (*Eretmochelys imbricata*) nesting at Bahía de Jiquilisco, El Salvador, 2013 (n = 28).

| Parameter | Mean | 95% CI |  |
| --- | --- | --- | --- |
|  |  | Low | High |
| Arsenic | 0.245 | 0.100 | 0.390 |
| Lead | 0.045 | 0.038 | 0.056 |
| Mercury | 0.008 | 0.004 | 0.017 |

## Discussion

Our results provide the first assessment of hematology, biochemistry, heavy metal, and persistent organic pollutant levels in the blood of wild hawksbills nesting in mangrove estuaries and establish baseline values for mature female hawksbills in these habitats in the eastern Pacific Ocean. The population sampled in this study was rated overall as healthy, as nesting hawksbills were in good body condition, had minimal epibiota, and generally had normal physical exam findings. While interpreting the parameters in this study, it is important to note that it is common that highly contaminated reptiles show no acute signs of health distress, thus our results should not be misinterpreted as confirming the species is healthy in the region.

### Hematology and Biochemistry

The hematological and biochemistry results are generally comparable to those of other species of sea turtles sampled with healthy populations (42) (43) (44) (45) including hawksbill nesting females at open-coast beaches in Brazil (1) and for hawksbill foraging aggregations at coral reefs in the eastern Pacific (Table 4). Some differences are notable in comparing values between studies, for example glucose in the (3) study is significantly higher than that of all other studies and the Packed Cell Volume is lower in our study than in (1). Notably only eight individuals were sampled in the (3) study and the sea turtles were caught in the open water and brought on to the beach instead of testing nesting females. The authors speculate that handling stress induced a stress hyperglycemia. This study also is the only other adult wild hawksbill study to include a white blood cell differential count, which varies from ours in numbers of heterophils and lymphocytes. The other two studies, (2) and (1) have similar values for biochemistries to our study.

**Table 4.** Available blood values for hawksbill turtles.

| Parameter<sup>a</sup> | Wrobel Goldberg et al. 2013 |  |  | Tobón-López & Amorocho Llanos 2014 |  |  | Muñoz-Pérez et al. 2017 |  |  | Tauer et al. this study |  |  |
| --- | --- | --- | --- | --- | --- | --- | --- | --- | --- | --- | --- | --- |
|  | Mean | SD | n<sup>b</sup> | Mean | SD | n<sup>c</sup> | Mean | SD | n<sup>d</sup> | Mean | SD | n<sup>e</sup> |
| *Hematology* |  |  |  |  |  |  |  |  |  |  |  |  |
| PCV (%) | 39.4 | 2.9 | 41 | — | — | — | — | — | — | 28.34 | 7.71 | 28 |
| RBC ($\times 10^{12}/L$) | — | — | — | — | — | — | 0.35 | 0.09 | 8 | — | — | — |
| WBC ($\times 10^9/L$) | — | — | — | — | — | — | 5.31 | 3.86 | 8 | — | — | — |
| Heterophils (%) | — | — | — | — | — | — | 32.3 | 6.9 | 8 | 65.5 | 12.81 | 28 |
| Lymphocytes (%) | — | — | — | — | — | — | 45.9 | 6.1 | 8 | 22.21 | 10.4 | 28 |
| Monocytes (%) | — | — | — | — | — | — | 3.6 | 1.9 | 8 | 6.04 | 2.92 | 28 |
| Basophils (%) | — | — | — | — | — | — | 0.1 | 0.2 | 8 | 2.82 | 2.20 | 28 |
| Eosinophils (%) | — | — | — | — | — | — | 18.5 | 4.5 | 8 | 0.36 | 0.99 | 28 |
| *Biochemistry* |  |  |  |  |  |  |  |  |  |  |  |  |
| Glucose (mg/dL) | 98.6 | 14.6 | 41 | 103.5 | 16.6 | 11 | 1567.6 | 180.2 | 7 | 99.71 | 21.56 | 51 |
| AST (U/L) | 55.4 | 7.1 | 41 | 132.6 | 111.2 | 11 | 196 | 54 | 8 | 46.33 | 14.29 | 51 |
| ALT (U/L) | 6.6 | 2.4 | 41 | — | — | — | 38 | 15 | 8 | 47.18 | 15.75 | 33 |
| ALP (U/L) | 15.9 | 3.7 | 41 | — | — | — | 53 | 26 | 8 | 56.63 | 17.33 | 51 |
| LDH (U/L) | — | — | — | 136.5 | 78.9 | 11 | — | — | — | 394.41 | 288.12 | 51 |
| Cholesterol (mg/dL) | 287 | 42 | 41 | 84.5 | 30.9 | 11 | — | — | — | 236.92 | 66.24 | 51 |
| TP (g/dL) | 5.45 | 0.63 | 41 | 2.5 | 0.7 | 11 | 4.8 | 0.7 | 8 | 3.79 | 0.62 | 51 |
| Phosphorus (mg/dL) | 11.3 | 1.4 | 41 | 6.7 | 2 | 11 | — | — | — | 9.76 | 3.23 | 18 |
| Calcium (mg/dL) | 11.6 | 1.5 | 41 | 7.8 | 1.8 | 11 | — | — | — | 11.09 | 5.14 | 36 |
| Sodium (mmol/L) | 139.6 | 3.5 | 41 | — | — | — | 157 | 2 | 7 | 155.04 | 7.33 | 51 |
| Potassium (mmol/L) | 5.09 | 0.76 | 41 | — | — | — | 4.2 | 0.4 | 7 | 4.68 | 0.51 | 18 |
| Chloride (mmol/L) | — | — | — | — | — | — | — | — | — | 122.71 | 13.14 | 51 |
| Biocarbonate (mmol/L) | — | — | — | — | — | — | — | — | — | 21.61 | 6.05 | 18 |
| Uric Acid (mg/dL) | 0.95 | 0.17 | 41 | 3.7 | 2.7 | 11 | — | — | — | 1.39 | 0.20 | 33 |
| Anion Gap (mmol/L) | — | — | — | — | — | — | — | — | — | 22.22 | 7.45 | 18 |

<sup>a</sup>PCV, packed cell volume; RBC, red blood cells; WBC, white blood cells; AST, aspartate aminotransferase; ALT, alanine aminotransferase; ALP, alkaline phosphatase; CK, creatine kinase; LDH, lactate dehydrogenase; TP, total protein.
<sup>b</sup>Hawksbills nesting on open-coast beaches in Brazil, Atlantic Ocean.
<sup>c</sup>Hawksbills foraging at coral reefs in Colombia, Pacific Ocean.
<sup>d</sup>Hawksbills foraging at coral reefs in Ecuador, Pacific Ocean.
<sup>e</sup>Hawksbills nesting on beaches within mangrove estuaries in El Salvador, Pacific Ocean.

One female in our study had white blood cell count and heterophil count twice as high as the lowest WBC and heterophil count sampled, so occult illness in one or more individuals of our studied population may be possible (46). Biochemistry reference ranges were established (2) for juvenile hawksbills occupying a coral reef ecosystem off the Pacific coast of Colombia, with calcium, total protein, phosphorus, glucose values similar to our data, but with much wider ranges of LDH, AST, and cholesterol. Some differences were noted between several hematological and biochemistry values when compared to published data from juvenile hawksbills undergoing rehabilitation in the United Arab Emirates (47). For example, juvenile hawksbills had lower mean PCVs, lower total white blood cell counts, and higher AST, CK, and uric acid levels. Additionally, mean calcium, phosphorus, and total protein levels were lower in the rehabilitated animals when compared to our study sample. These differences may be due, at least in part, to the impaired health of animals in rehabilitation, as well as possible geographic variation in environmental variables or in the life-history characteristics of hawksbills in distinct oceanic regions. Variation in biochemistry ranges may reflect differences in physiological requirements between life stages (i.e. juvenile vs. adults) and/or behavior/habitats (nesting in mangrove estuaries vs foraging at coral reef ecosystems) of each studied hawksbill population (15).

### Heavy Metals

Heavy metal values appear variable amongst species and subpopultations in loggerhead (*Caretta caretta),* kemp’s ridley (*Lepidochelys kempii),* and green (*Chelonia mydas*) turtles (48) (49) (50), and are likely related to environmental effects, diet, age, and geography. While hawksbills are omnivorous, their diet worldwide is primarily composed of sponges (51), which are of low trophic level and may explain lower levels of contaminants than sea turtle species that eat items higher up on the food chain, such as olive ridley (*Lepidochelys olivacea*) and kemp’s ridley turtles (48). Adult hawksbills have been documented having relatively low concentrations of the heavy metals in their blood, although maternal transfer of heavy metals from adult hawksbills to their eggs is known to occur (43).

Higher levels of arsenic were found in adult hawksbill tissues in Japan, particularly in muscle, than compared to adult green turtles (52). Additionally, arsenic levels of marine sponges were found to range from 0.8–157 mgm/gram of dry weight, suggesting that sponges may be a significant source of arsenic in adult hawksbills. It is unclear the role that sponges may play in accumulation of other heavy metals or persistent organic pollutants, such as the low levels of lead and mercury found in our study population.. Importantly, hawksbills in our study area utilize mangrove estuaries and are believed to feed predominantly on mangrove seeds and roots (M. Liles, pers. obs.), indicating that they may feed at an even lower trophic level than populations of hawksbills in other regions. The tendency to feed at low trophic levels may enable eastern Pacific hawksbills to avoid higher levels of blood pollutants seen in conspecifics in other habitats, as well as other sea turtle species.

### Persistent Organic Pollutants

Organic and inorganic pollutants have been more frequently studied in loggerheads than other sea turtle species (53) (54) (55). Studies on loggerheads have found detectable POP and PCB results in which several of the individual contaminants had correlations with changes in clinical parameters such as packed cell volume (56). Further studies are needed on all sea turtle species to determine the individual and population level effects on health and reproductive outcomes in animals exposed to inorganic and organic pollutants.

The trophic level of food items consumed by sea turtle species at different life stages may impact levels of POP and PCBs. For instance, green turtles consume marine invertebrates as juveniles before transitioning to primarily algae and sea grass as adults, whereas adult leatherback and hawksbills forage on jellyfish and primarily marine sponges, respectively (57). For hawksbills and leatherbacks, this may mean they tend to accumulate more polluntants. More recently, however, leatherback turtles (*Dermochelys coriacea*) in Gabon with evaluated levels of POP and PCB in the blood of nesting and all turtles had levels below the detectable limit (42), a recent study (58) noff the west coast of Senegal in the Cape Verde Islands comparing POP and PCB levels in juvenile green and hawksbill turtles found detectable levels in both species, although green turtles had both higher levels and a greater prevalence of contamination. Trophic levels might not reflect higher levels of POP in adult green and hawksbill turtles and viable turtle eggs (59).

## Conclusions

Our study provides baseline health data for hawksbills nesting at a primary rookery located in a mangrove estuary in the eastern Pacific Ocean, which can provide a starting point for long-term monitoring of health status of hawksbills in the region and offer diagnostic indications for treatment of individuals in clinical rehabilitation. Additional studies between healthy juvenile and adult hawksbills in both mangrove estuaries and other habitats should be conducted to delineate size or age related differences in biochemistry and hematologic values in this species, as apparent health status may not reflect contaminant loads. We suggest that future research determine contaminant loads of marine sponges and mangrove vegetation in the Bahía de Jiquilisco and the potential role they play in accumulation of toxins in the environment. It is possible ecosystem processes are occurring that prevent uptake of toxins in the environment to the sea turtles themselves, or through their diet, which, contrary to most hawksbill populations, includes substantially more vegetation (60). Further studies at Bahia de Jiquilisco utilizing skin, muscle, carapace, fat and liver may provide different results than those obtained in this study.

## Acknowledgements

We thank local egg collectors at Bahía de Jiquilisco for their collaboration—this study would have been impossible without their knowledge and participation. We acknowledge N. Sanchez, O. Rivera, F. López, G. Serrano Liles, D. Melero, E. LaCasella, F. Rivas, M. Hidalgo, W. J. Nichols, J. Urteaga, E. Ramírez, and C. Dueñas for assistance. We are grateful to the following institutions for permits: Ministry of Environment and Natural Resources in El Salvador (sample collection: MARN-AIMA-DGBPN-GVS-054-2013; MARN-DEVS-GVS-26-2014), Ministry of Agriculture and Ranching in El Salvador (sample export: CITES 09105), and Southwest Fisheries Science Center (NMFS-NOAA) in United States (sample import: CITES 13US844694/9). Research ethics were evaluated by Cūra Earth’s scientific advisory group. We thank The Rufford Foundation for funding.

